# Cell-free DNA (cfDNA) and exosome profiling from a year-long human spaceflight reveals circulating biomarkers

**DOI:** 10.1101/2020.11.08.373530

**Authors:** Daniela Bezdan, Kirill Grigorev, Cem Meydan, Fanny A. Pelissier Vatter, Michele Cioffi, Varsha Rao, Kiichi Nakahira, Philip Burnham, Ebrahim Afshinnekoo, Craig Westover, Daniel Butler, Chris Moszary, Matthew MacKay, Jonathan Foox, Tejaswini Mishra, Serena Lucotti, Brinda K. Rana, Ari M. Melnick, Haiying Zhang, Irina Matei, David Kelsen, Kenneth Yu, David C Lyden, Lynn Taylor, Susan M Bailey, Michael P.Snyder, Francine E. Garrett-Bakelman, Stephan Ossowski, Iwijn De Vlaminck, Christopher E. Mason

**Affiliations:** Department of Physiology and Biophysics, Weill Cornell Medicine, New York, NY, USA; Children’s Cancer and Blood Foundation Laboratories, Departments of Pediatrics, and Cell and Developmental Biology, Drukier Institute for Children’s Health, Meyer Cancer Center, Weill Cornell Medical College, New York, NY, USA; Department of Genetics, Stanford University School of Medicine, Stanford, CA, USA; Nara Medical University, Kashihara, Nara, Japan; Department of Bioengineering, University of Pennsylvania, Philadelphia, PA 19104; The HRH Prince Alwaleed Bin Talal Bin Abdulaziz Alsaud Institute for Computational Biomedicine, Weill Cornell Medicine, New York, NY, USA; The WorldQuant Initiative for Quantitative Prediction, Weill Cornell Medicine, New York, NY, USA; Department of Psychiatry University of California, San Diego, La Jolla, CA, USA; Department of Medicine, Weill Cornell Medicine, New York, NY, USA; Department of Medicine, Memorial Sloan Kettering Cancer Center, New York, NY, USA; Department of Environmental & Radiological Health Sciences, Colorado State University, Fort Collins, CO, USA; Department of Medicine, University of Virginia School of Medicine, Charlottesville, VA, USA; Department of Biochemistry and Molecular Genetics, University of Virginia School of Medicine, Charlottesville, VA, USA; University of Virginia Cancer Center, Charlottesville, VA; Institute of Medical Genetics and Applied Genomics, University of Tübingen, Tübingen, Germany; Nancy E. and Peter C. Meinig School of Biomedical Engineering, Cornell University, Ithaca, NY, USA; The Feil Family Brain and Mind Research Institute, Weill Cornell Medicine, New York, NY, USA

**Keywords:** NASA, cfDNA, liquid biopsy, mtDNA, mtRNA, exosomes, International Space Station, NASA Twins Study

## Abstract

The health impact of prolonged space flight on the human body is not well understood. Liquid biopsies based on cell-free DNA (cfDNA) or exosome analysis provide a noninvasive approach to monitor the dynamics of genomic, epigenomic and proteomic biomarkers, and the occurrence of DNA damage, physiological stress, and immune responses. To study the molecular consequences of spaceflight we profiled cfDNA isolated from plasma of an astronaut (TW) during a year-long mission on the International Space Station (ISS), sampling before, during, and after spaceflight, and compared the results to cfDNA profiling of the subject’s identical twin (HR) who remained on Earth, as well as healthy donors. We characterized cfDNA concentration and fragment size, and the positioning of nucleosomes on cfDNA, observing a significant increase in the proportion of cell-free mitochondrial DNA inflight, suggesting that cf-mtDNA is a potential biomarker for space flight-associated stress, and that this result was robust to ambient transit from the International Space Station (ISS). Analysis of exosomes isolated from post-flight plasma revealed a 30-fold increase in circulating exosomes and distinct exosomal protein cargo, including brain-derived peptides, in TW compared to HR and all known controls. This study provides the first longitudinal analysis of astronaut cfDNA during spaceflight, as well as the first exosome profiles, and highlights cf-mtDNA levels as a potential biomarker for physiological stress or immune system responses related to microgravity, radiation exposure, and other unique environmental conditions on the ISS.

## Introduction

A wide range of physiological effects impact the human body during a prolonged stay in microgravity, such as headward fluid shift, atrophy of muscles, and decreases in bone density, which have been described for astronauts on the international space station (ISS)(Williams et al., 2009). In recent years, an increasing number of government and private space agencies have formed, and missions to the Moon and Mars are now planned for the late 2020s and 2030s (Iosim et al, 2020). These pending missions may span 30 months and require landing on a planet with almost no clinical infrastructure for medical monitoring or treatments. Yet, data on physiological changes of long-term missions (>6 months) is almost non-existent. These long-duration missions and the increasing exposure of humans to spaceflight-specific conditions necessitates the study of molecular changes in the human body induced by exposure to spaceflight stressors such as microgravity, radiation, noise, restricted diet, and reduced physical work opportunities. The NASA Twins study (Garrett-Bakelman et al., 2019) enabled interrogation of the impact of prolonged spaceflight on the human biology and cell-to-cell variations in the immune system (Gertz et al., 2020); however, there has never been a study on the impact of spaceflight on cell-free DNA (cfDNA).

Molecular signatures informative of human health and disease can be found in cfDNA and nucleic acids isolated from plasma, saliva, or urine (Heitzer et al., 2018; Hummel et al., 2018; Siravegna et al., 2017; Verhoeven et al., 2018; Volik et al., 2016). Non-invasive methods for monitoring health-related biomarkers in liquids such as plasma (‘liquid biopsy’) have already been successfully introduced in a wide range of contexts, including: prenatal testing for detection of trisomy and micro-deletions (Bianchi et al., 2014; Zhang et al., 2019), cancer diagnostics (Bettegowda et al., 2014; Diehl et al., 2008; Wang et al., 2017), monitoring of cancer therapies (Birkenkamp-Demtröder et al., 2016; Wan et al., 2020), monitoring of the health of solid-organ transplants (Verhoeven et al., 2018; De Vlaminck et al., 2014), and screening for infections (Blauwkamp et al., 2019; Burnham et al., 2018; De Vlaminck et al., 2013). Hence, liquid biopsy is a potentially useful method for monitoring physiologic conditions of astronauts before, during and after spaceflight.

Indeed, cfDNA is extremely dynamic and responsive, providing strong indicators of DNA damage and tumor growth in distal tissues (Newman et al., 2016), immune response or infection (Zwirner et al., 2018), and RNA regulatory changes, with an innate capacity to reveal the cells of origin undergoing apoptosis or necrosis (Thierry et al., 2016). Various studies have reported changes in cfDNA concentration (Zwirner et al., 2018), cfDNA fragment length distribution (Mouliere et al., 2011; Underhill et al., 2016), mutation profiles and signatures (Newman et al., 2016), and cfDNA methylation (Shen et al., 2018) indicative of physiological conditions such as cancer. Mitochondrial DNA (mtDNA) can also be found in the extracellular space, circulating as short DNA fragments, encapsulated in vesicles and even as whole functional mitochondria (Amir Dache et al., 2020; Song et al, 2020). Several recent studies observed increased levels of cell-free mitochondrial DNA (cf-mtDNA) in psychological conditions (Lindqvist et al., 2016, 2018) and reduced cf-mtDNA levels in Hepatitis B infected patients associated with a higher risk of developing hepatocellular carcinoma (Li et al., 2016). However, since no such information exists for using these metrics for astronauts, we investigated the utility of cfDNA for the monitoring of the physiologic conditions of astronauts to spaceflight.

Of note, cfDNA comprises the footprints of nucleosomes, and these nucleosome features enable tracing of the tissue-of-origin for cfDNA in normal and disease states, through analysis of nuclear architecture, gene structure and expression (Murtaza and Caldas, 2016; Snyder et al., 2016). In particular, nucleosome positioning and depletion of short cfDNA sequences reveal footprints of transcription factor binding, promoter activity, and splicing, ultimately informing gene regulatory processes in the tissue/cell of origin (Snyder et al., 2016). Similar information can be revealed from exosomes, which are nano-sized vesicles (size 30–150nm) derived from perinuclear luminal membranes of late endosomes/multivesicular bodies and released into extracellular environment via multivesicular body fusion within the cell membrane ((Kalluri and LeBleu, 2020; Mathieu et al., 2019)) that can mediate long-range physiological crosstalk (Hoshino et al., 2015; Mathieu et al., 2019). Exosomes act as vehicles for horizontal transfer of information through their cargo: proteins, lipids, metabolites and DNA, as well as coding and non-coding RNAs (Valadi et al., 2007; Wortzel et al., 2019). Moreover, exosomes can be powerful mediators of responses to environmental stimuli as external and physiological stress impact their release, cargo and function, contributing to pathogenesis (Harmati et al., 2019; O’Neill et al., 2019; Qin et al., 2020). Since exosomes are abundant in plasma, they are critical components of liquid biopsies (Colombo et al., 2014; Hoshino et al., 2020) and analysis of their content can complement the information obtained from cfDNA, but there is no information about exosomes in astronauts.

To address this gap in knowledge, we profiled cfDNA isolated from plasma samples before, during, and after the one-year mission on the International Space Station (ISS) to evaluate the utility of cfDNA as a means to monitor physiological problems during extended missions in space. We also profiled the exosomes of both astronauts after the mission completion. While bulk RNA sequencing data have shown widespread gene expression changes in astronauts, including mitochondrial RNA (mtRNA) spikes in flight samples from the One-Year Mission (Garrett-Bakelman et al., 2019), there has not yet been a study of astronauts that has leveraged cfDNA and exosomes. We focused on quantitative measures such as the levels of mitochondrial DNA, cfDNA fragment length, and the depletion of nucleosome signatures at transcription start sites. Together, our NGS results provide a “whole-body molecular scan”, which can provide a novel measurement of the impact of spaceflight on the human body, as well as serve as a continued metric of physiology and cellular stress for future long-during missions.

## Results

### Study design and sample collection

We analyzed circulating cfDNA of a pair of male monozygotic twins over two years, starting when they were both 50 years old. During the NASA Twin Study, the flight subject (TW) was aboard the International Space Station (ISS) for 340 days, while his identical twin, the ground subject (HR), remained on Earth. We collected cfDNA at 12 time points from HR and 11 time points from TW. Of the latter, four samples were collected inflight on board of the ISS or space shuttle. In addition, we profiled the cfDNA of an unrelated control subject (MS) to simulate the ambient return from the ISS. To control for ambient return (AR) effects (return of samples in the Soyuz capsule) on the molecular signatures of cfDNA, we subjected two MS samples and one HR sample to an extended shipping procedure (see Methods). Plasma and cfDNA were extracted using the same protocol for all samples (Methods). We observed a broad range of cfDNA concentration between 6.7 ng/ml and 79.9 ng/ml plasma (mean = 27.9 ng/ml, median = 23 ng/ml) across samples (**Table 1**). However, we found no significant difference in cfDNA concentrations between flight, ground or control subjects (ANOVA p = 0.49, **Supp. Fig. 1A**), TW and HR (Wilcoxon rank test p = 0.65), and flight and ground samples (Wilcoxon rank test p = 0.352). TW showed borderline significantly higher cfDNA concentration pre- and post-flight compared to inflight (Wilcoxon rank test p = 0.043), however, this is not significant when comparing TW inflight, TW ground, and HR/MS ground (ANOVAR p = 0.4, **Supp. Fig 1B**). Complementary metadata on the health status of TW and HW during the mission has been previously published (Garret-Bakelman et al., 2019), and no deviations in medication or exercise regimen were noted in the medical records.

**Table 1.**
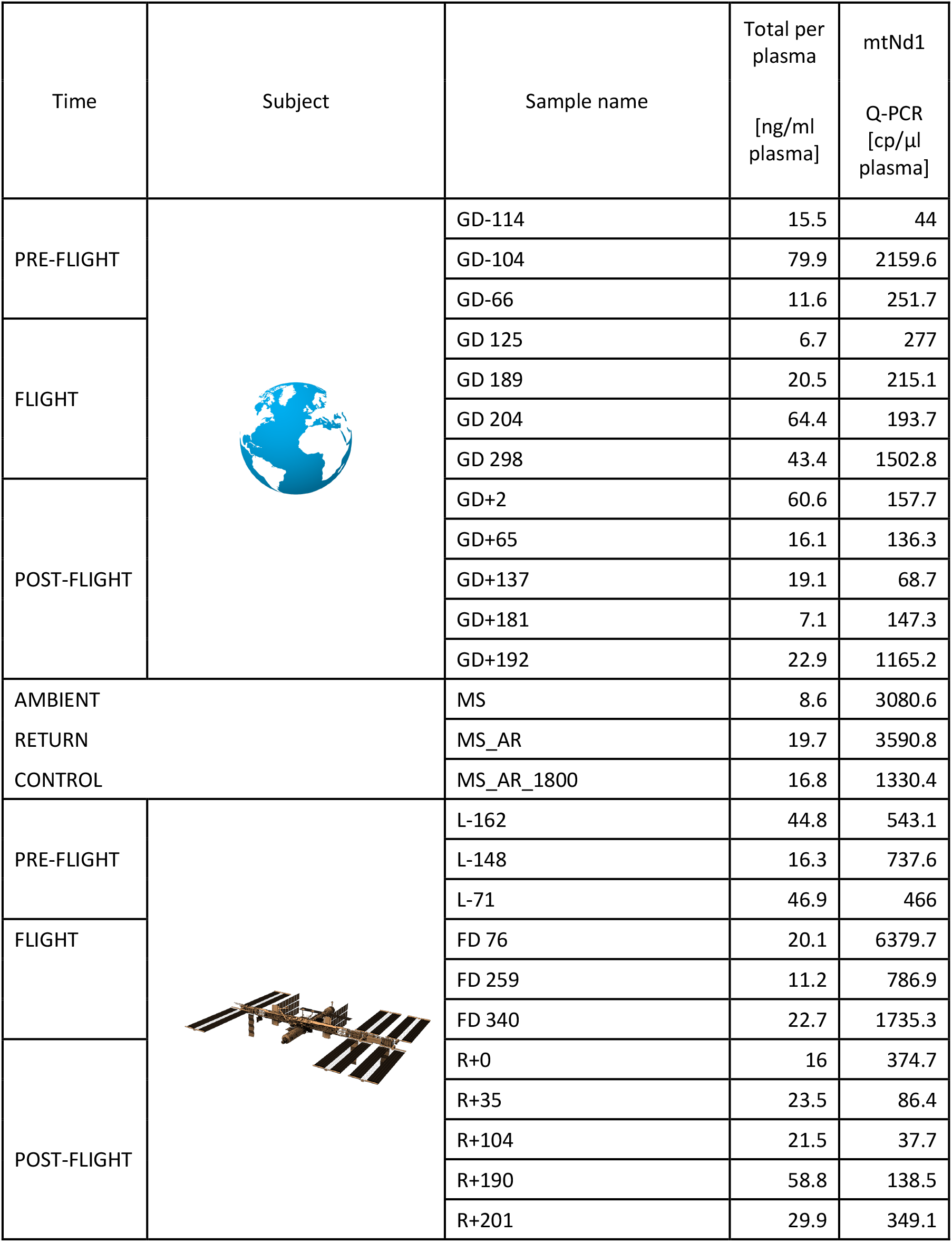
Overview of all plasma samples obtained during the 1-year mission. Subjects for this mission included the ground subject HR (blue), flight subject TW (green) and control subject MS (yellow). Samples taken on the ISS are highlighted in red. The last two columns show the concentration of cfDNA per ml plasma and the Q-PCR results for the mitochondrial transcript mtNd1 in copy/µl plasma.

### Cf-DNA fragment length distribution is influenced by the ambient return

It has previously been shown that cfDNA derived from tumor cells is shorter than cfDNA derived from healthy cells (Jiang et al., 2015; Mouliere et al., 2011). This effect can be explained by a change in nucleosome binding or by a degradation of nucleotides at the end of nucleosome loops. We therefore hypothesized that environmental stressors such as microgravity or radiation could also impact the length distribution of cfDNA. Indeed, we found a slight shift to longer cfDNA fragment lengths in TW inflight samples (**Fig. 1A**). However, a similar shift was observed in ground samples subjected to ambient return simulation (**Fig. 1A**, boxplots with yellow border). Ambient return samples show a similar peak at the 300 to 400bp fragment length, which is only marginally visible for fresh samples (**Fig. 1B**). Thus, some proportion of long cfDNA fragments likely originate from blood cells damaged during return flight or transport from the ISS.

**Figure 1.**
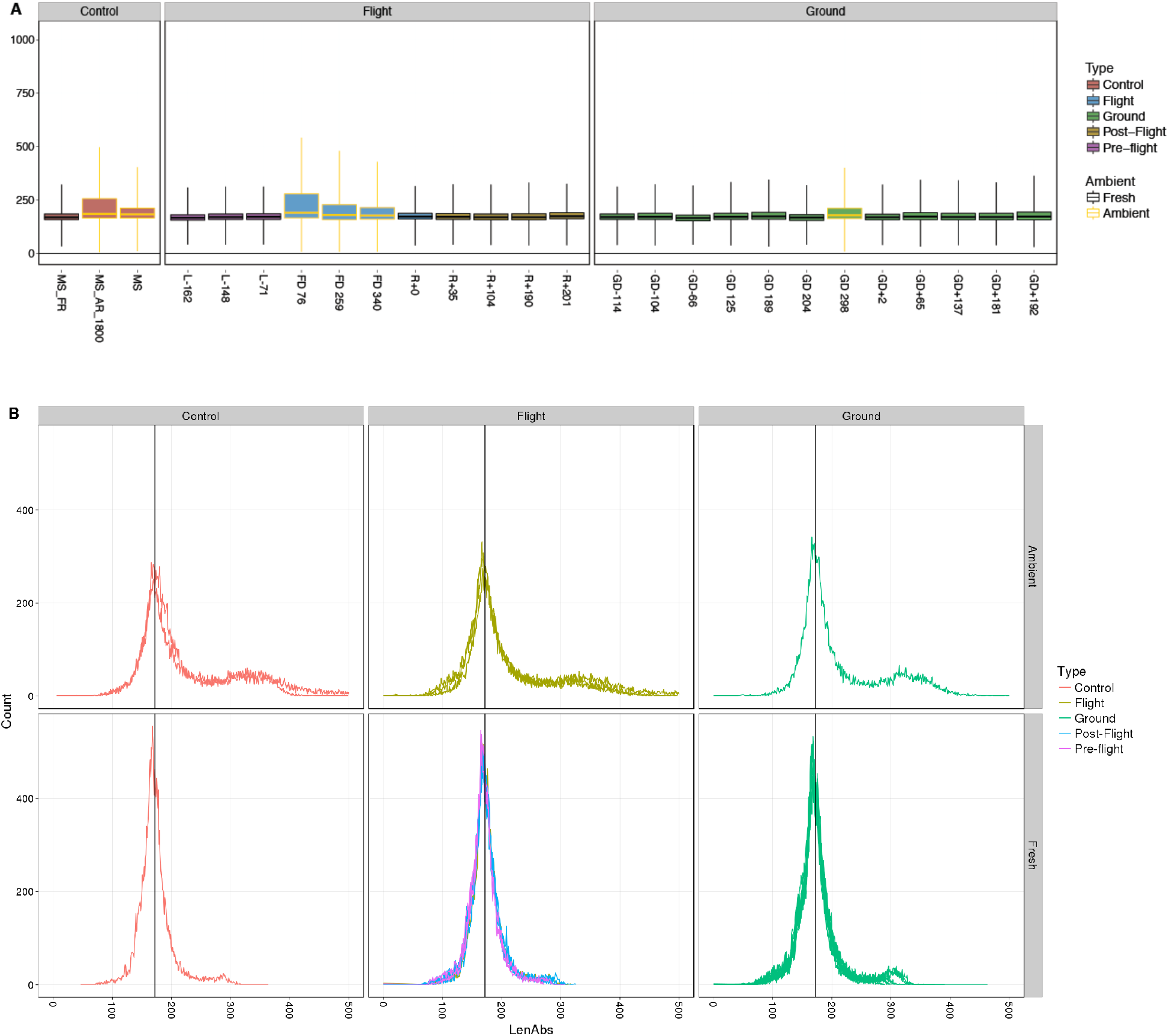
Size distribution of cfDNAs in ambient return, ambient return simulation and fresh samples. (A). Ambient return simulation samples (control and ground samples with yellow border) show a highly similar pattern as observed for inflight samples (blue box with yellow border). Long cfDNA fragments likely originate from blood cells damaged during transport. (B) Ambient return samples show an increased fraction of cfDNA with fragment length > 300bp compared to fresh samples. Our experimental procedure does only allow interrogation of DNA fragments up to a length of 500bp, thus the content of long mtDNA fragments contained in intact circulating mitochondria is not reflected in this analysis.

To examine how this might affect other cfDNA fractions, we next examined cell-free mitochondrial DNA (cf-mtDNA). Recent studies indicate that a prominent fraction of cf-mtDNA in the plasma is contained within intact, circulating mitochondria (Al Amir Dache, 2020) and that larger mtDNA fragments can also arise from blood cell degradation. However, our centrifugation step largely removed intact mitochondria and our library preparation comprised mostly smaller DNA fragments (at least 75% are <350bp)(**Supplemental Fig. 2**), including an even smaller fraction (<10%) of the aligned reads (**Fig. 2**). Thus, the observed fractions of cf-mtDNA are mostly derived from shorter cf-mtDNA molecules and should represent cf-mtDNA that is randomly fragmented and sequenced across the entire mitochondria.

**Figure 2.**
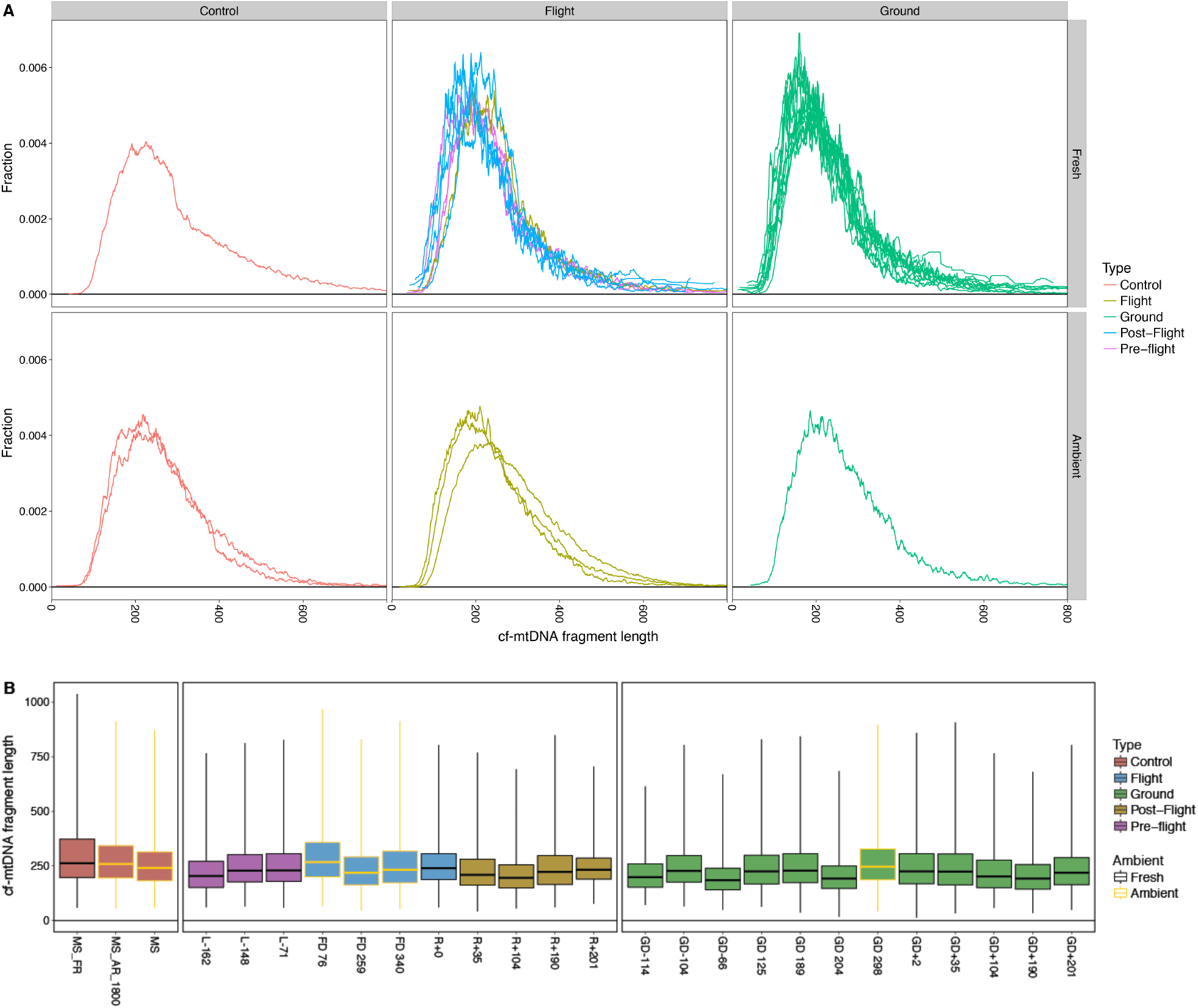
Size distribution of cf-mtDNAs. We observed a wider range of cf-mtDNA lengths compared to total cfDNA (from 100 to 600bp). (A) cf-mtDNA size distributions are similar in ground, flight and control samples, and are not affected by ambient return (AR) or AR simulation. (B) Average length of cf-mtDNA is significantly longer than the average length reported for chromosomal cfDNA (∼250bp vs. ∼160bp). The average length of cf-mtDNA is not affected by sample type (control, flight, ground) or sample handling (fresh, AR, AR simulation).

As further evidence of this, the mitochondrial genome showed continuous read coverage in all samples, ranging from 50x-200x coverage (**Supplemental Fig. 3**), regardless of the collection method. Indeed, the length distribution of cf-mtDNA is not affected by ambient return as observed for chromosomal cfDNA (**Fig. 2B**), and the average length does not change significantly in inflight samples or AR simulation samples. Even though cf-mtDNA amounts can significantly vary based on the donor profiles (Lindqvist et al, 2016) and degree intact vs. fragmented mitochondria, these NGS data showed that the total cf-mtDNA profiles show relative uniformity in both length and proportion of reads (**Fig. 2**).

### Levels of cell-free mitochondrial DNA are increased during space flight

Next we investigated the fraction of cf-mtDNA relative to chromosomal cfDNA in plasma of TW, HR, and MS. In order to characterize the cfDNA originating from mitochondria during spaceflight, we normalized the count of NGS reads mapping to the mitochondrial chromosome (chrM) by chromosome length and the total number of reads in the library, generating a RPKM measurement. For comparison, we performed the same procedure with reads mapping to chromosome 21. We found a sharp increase of cf-mtDNA for subject TW for inflight samples (**Fig. 3A**) compared to TW ground samples (Wilcoxon rank test p= 0.012), compared to HR ground samples (Wilcoxon rank test p= 0.018), and compared to all ground samples of HR and TW (Wilcoxon rank test p=0.0045, ANOVA p=0.00049). In contrast, we found no significant increase in cfDNA mapping to chromosome 21 (**Fig. 3B**) in TW-inflight compared to ground samples of TW and HR.

**Figure 3.**
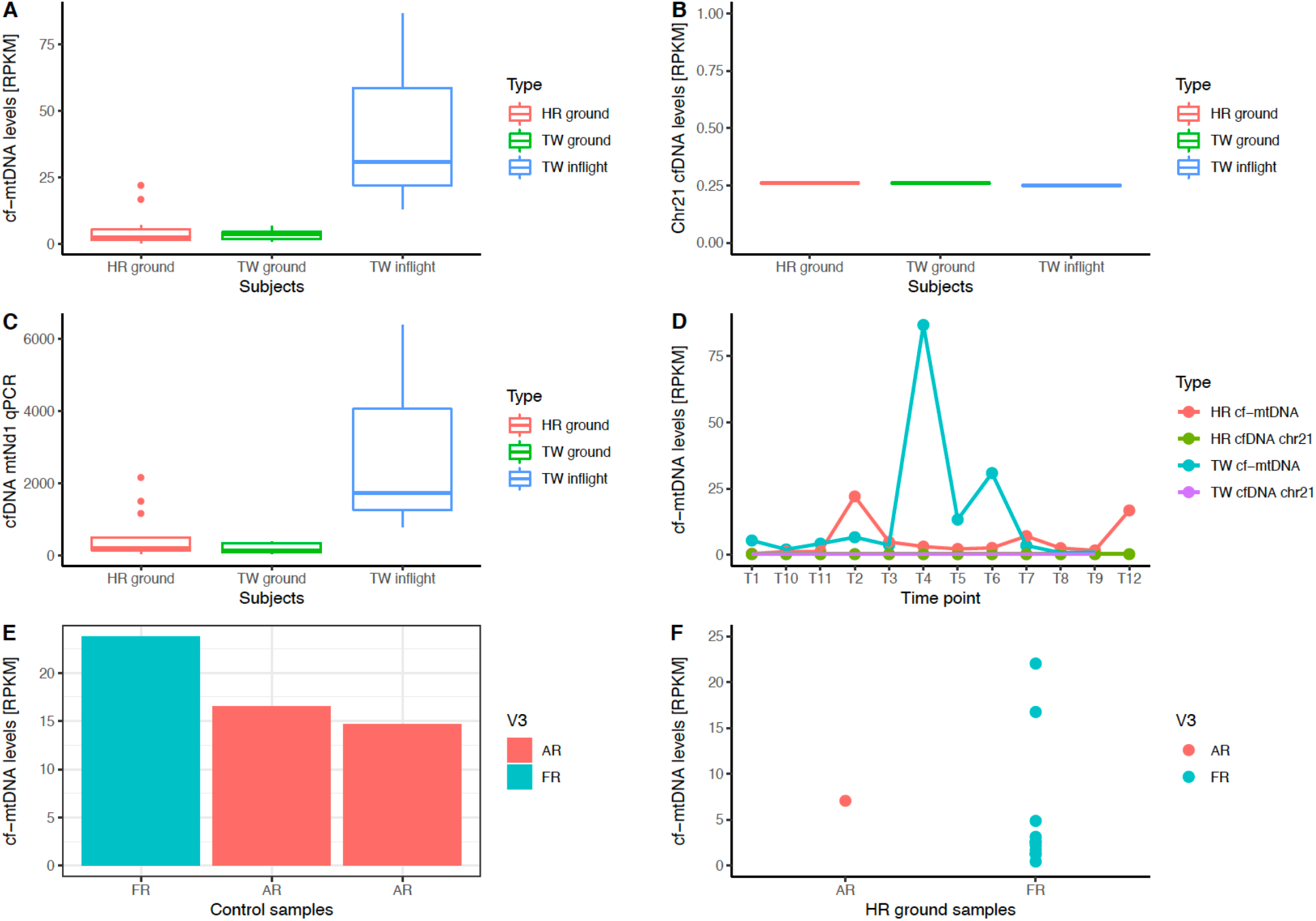
Analysis of normalized cfDNA read counts by chromosome, including the mitochondrial genome. (A) TW exhibits a significant increase in cell-free mtDNA during space flight compared to TW and HR ground samples. Counts are reads per kilobase per Million reads, or RPKM (B) Chromosomes do not show any change in RPKM during space flight, as exemplified using chr21. (C) Q-PCR based validation of increased cf-mtDNA fraction in plasma during space flight. (D) Normalized cf-mtDNA fraction and fraction of reads mapping to chr21 for 12 time point during the mission (T4-T6 = space flight). The highest increase in cf-mtDNA fraction is observed during the first months on ISS. (E) Ambient return simulation using two control samples showed no increase in cf-mtDNA compared to fresh samples, but a slight reduction. (F) Ambient return simulation (AR) using one HR ground sample did not show a significant increase in cf-mtDNA fraction. Two outliers within the fresh samples (FR) indicate that other conditions (e.g. stress, disease, immune reaction) could have influenced cf-mtDNA levels of HR on the ground.

Notably, the mtDNA levels in whole blood increased steadily inflight while on the ISS. Indeed, TW had the highest fraction of cf-mtDNA within the first inflight timepoint (T4), including more than a 24-fold increase, when compared to ground samples (**Fig. 3C, 3D**). In the two later inflight time points, he had 4- and 8-fold increases compared to pre-flight levels. The normalized levels of chromosome 21 cfDNA were stable for both TW and HR for the duration of the mission (0.25-0.26 RPKM), revealing no obvious bias due to sample handling (**Fig. 3D**). Interestingly, a positive correlation between mtDNA copy number and telomere length in healthy adults has been previously reported, and telomere elongation in blood and urine was also observed during spaceflight for TW (Garrett-Bakelman et al., 2019, Luxton et al, 2020).

Given the previously discussed effects of AR on cfDNA lengths, we tested for potential bias in cf-mtDNA levels due to AR. To do this, we compared the cf-mtDNA fraction observed in the MS simulated-AR samples (2 samples) to the MS control sample. We found that cf-mtDNA levels were actually lower in AR than in FR samples (**Fig. 3E)**, suggesting that the shipping procedure from the ISS is likely not causing the observed increase in cf-mtDNA levels seen in the inflight samples. In addition, the AR simulation of the ground subject (HR) did not show a significant increase of cf-mtDNA levels compared to other HR samples (**Fig. 3F**). Thus, these data suggest that the cf-mtDNA fraction was significantly increased during space flight, and not due to the AR blood-from-ISS transport process.

### Nucleosome positioning suggests a shift in cell of origin of cfDNA due to transport conditions

Given that nucleosome positions are associated with both cfDNA and gene expression (Jiang and Pugh, 2009), we computed the nucleosome depletion around nucleosomes at transcription start sites (TSS) to infer gene expression (**Fig. 4A**), as previously demonstrated by Ulz and colleagues (Ulz et al., 2016). Indeed, these data indicated that the strength of nucleosome depletion is correlated to bulk gene expression from RNA-seq of the same subjects (Garrett-Bakelman et al., 2019) (**Fig. 4A**), with a decreased coverage at the site of the transcriptional start site (TSS) for highly expressed genes. Second, we identified the nucleosome footprint of CTCF in gene bodies, hypothesizing that nucleosome positioning patterns could reveal broad changes in gene regulation during spaceflight. A t-SNE analysis of TW and HR samples showed no flight-specific clustering (**Fig 4B**), indicating that nucleosome positioning identified through cfDNA may not be sensitive enough to identify spaceflight-related gene expression changes.

**Figure 4.**
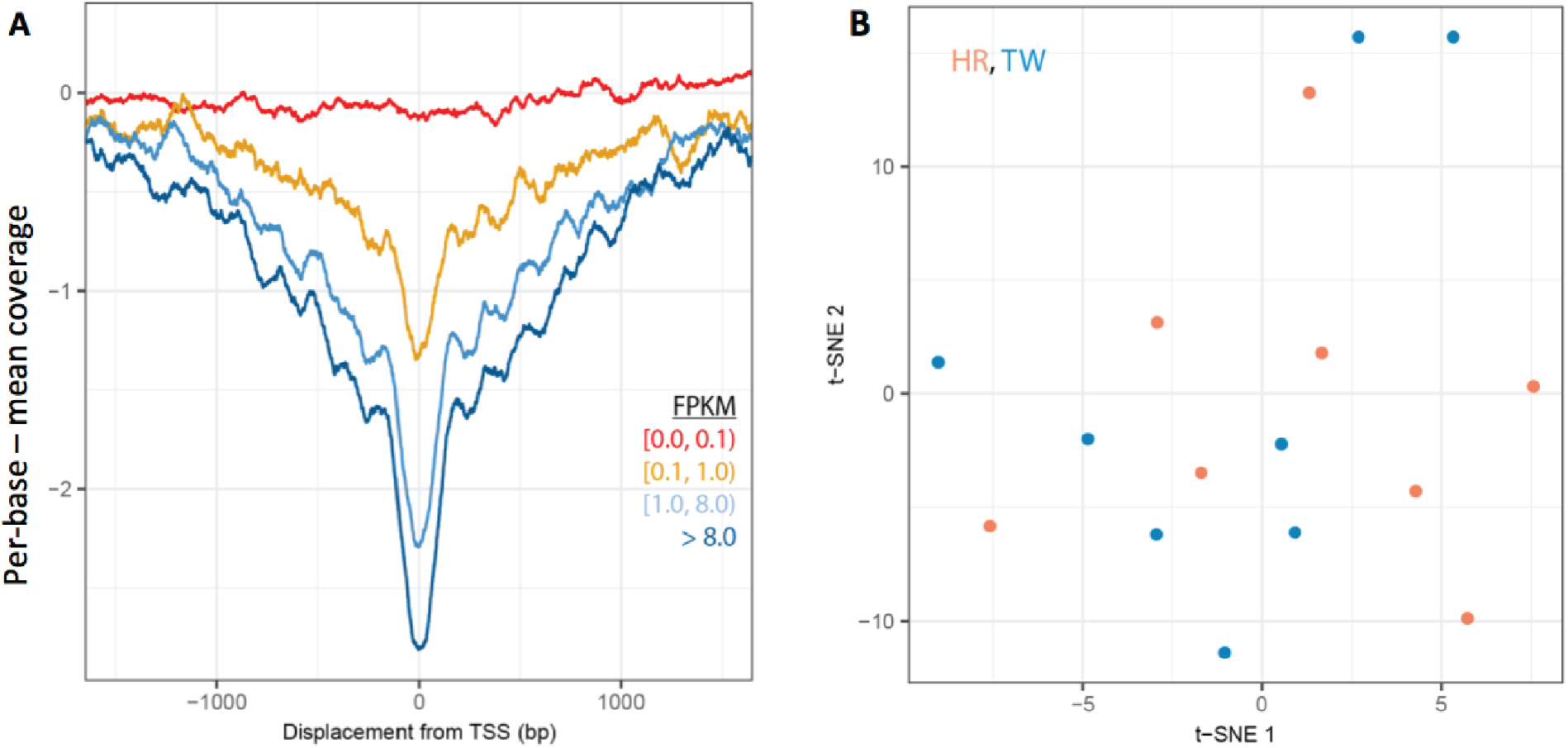
cfDNA nucleosome footprinting. (A) Nucleosome depletion in cfDNA around transcription start sites (TSS) is highly correlated with the expression of the respective genes and can therefore be used to estimate promoter activity and gene expression. (B) t-SNE based on genome-wide promoter nucleosome footprint of cfDNA samples reveals no clustering of flight subject and ground subject samples.

However, based on the correlations between per-tissue gene expression values (Kim et al., 2014) and nucleosome positioning observed on cfDNA, clear tissue signals in cfDNA were inferred for all plasma samples. Higher values (Pearson’s correlation coefficient) suggested higher gene expression and stronger tissue signal (**Fig. 5A**) for hematopoietic lineages (up to rho = 0.156, n = 1087411335), mid-range for liver, adrenal gland, and the retina (0.04-0.07) and less so for other peripheral tissues (e.g. lung, esophagus, 0.00-0.01). These results are consistent with the expected cfDNA prevalence in blood and with previous findings (Snyder et al., 2016). Despite such clear signals on tissue of origin, strong clustering of samples was observed, due to the confounding effect of ambient return. This was seen in both the tissue-of-origin analysis (**Fig 5A**) as well as TSS protection (**Fig. 5B**), highlighting the need for controls and correction for any degradation. Further, this analysis does not take into account the cf-mtDNA reads, and therefore may not reflect the tissue of origin for mitochondrial reads or heteroplasmy.

**Figure 5.**
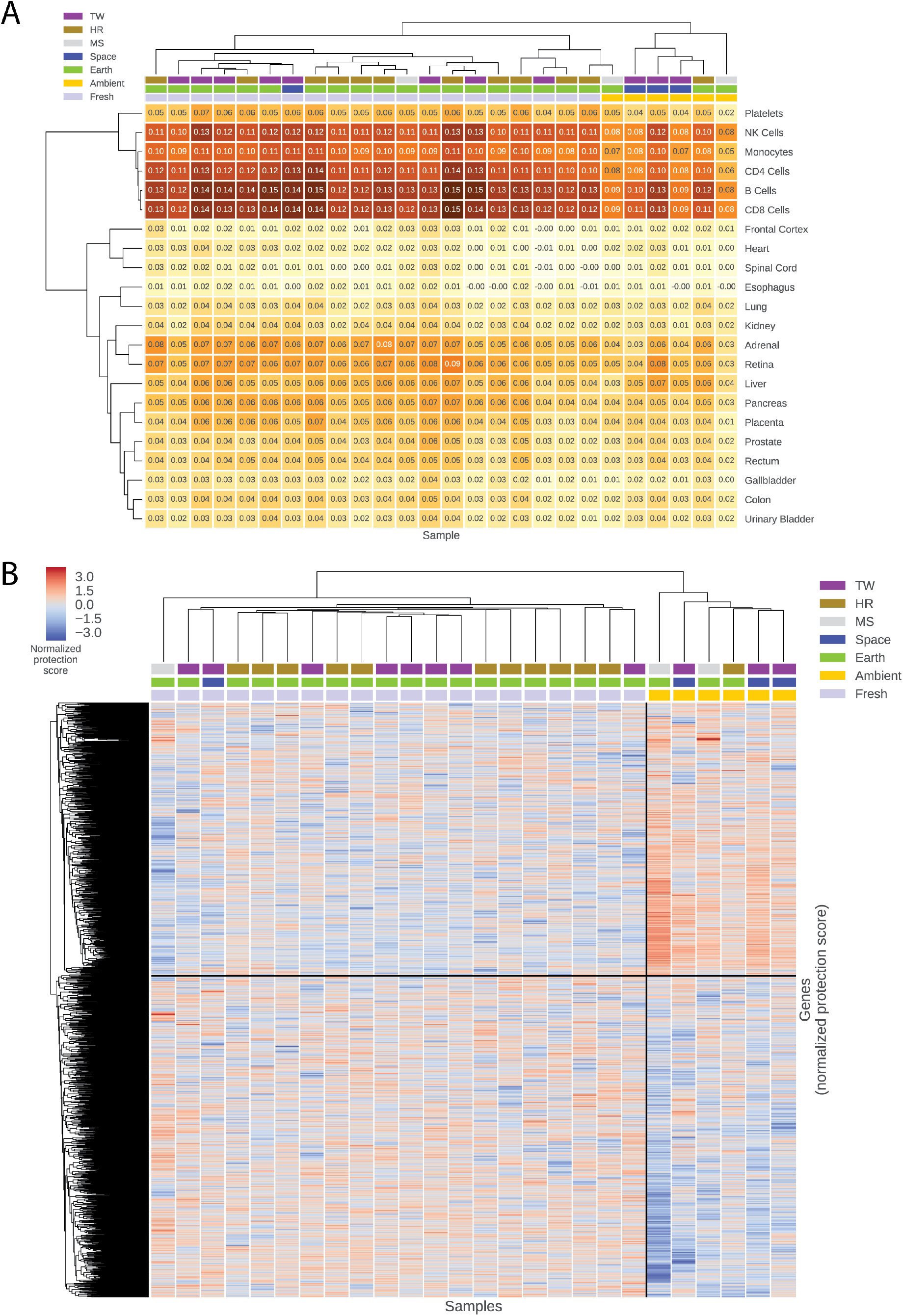
Tissue of origin deconvolution. **(A)** Correlation coefficients (multiplied by -1) for each tissue in each sample, clustered by sample and by tissue. The highest signals are, expectedly, from cells of hematopoietic origin. Spaceflight-dependent dynamics of tissue signal are confounded by the effect of ambient return, as suggested by ambient return samples tending to cluster together regardless of other features. (B) Clustering of samples using TSS protection in cfDNA as a measure of gene expression (lower protection correlates to higher expression). Ambient return samples cluster tightly together and uncover two major clusters of genes whose expression differs significantly from other samples, suggesting transport-related degradation processes or nucleosome detachment. Distribution of mean TSS protection per gene in ambient return and fresh samples is significantly different (t-test p<1e-3).

### Analysis of plasma-circulating exosomes post-flight

To determine how prolonged space missions and Earth re-entry impact circulating exosomes, we analyzed exosomes from the plasma of TW three years post-return to Earth, and compared their size, number and proteomes to plasma-derived exosomes isolated from HR and 6 age-matched, healthy controls. Exosomes were isolated by differential ultracentrifugation and both the size and number of exosomes were characterized by nanoparticle tracking analysis (NTA) (**Fig. 6A-E**). While the median size of exosomes was similar between HR, TW and healthy controls (**Fig. 6 A-D)**, the number of particles was ∼30 times higher in TW compared to HR and healthy controls (**Fig. 6 E**). Proteomic mass spectrometry analysis revealed that TW, HR and control exosomes packaged similar numbers of proteins, including a total of 191 exosomal proteins shared among all samples. HR’s exosome catalog contained 26 unique proteins, TW exosomes contained 61 unique proteins, and healthy controls contained 105 unique proteins (**Fig. 6F**).

**Figure 6.**
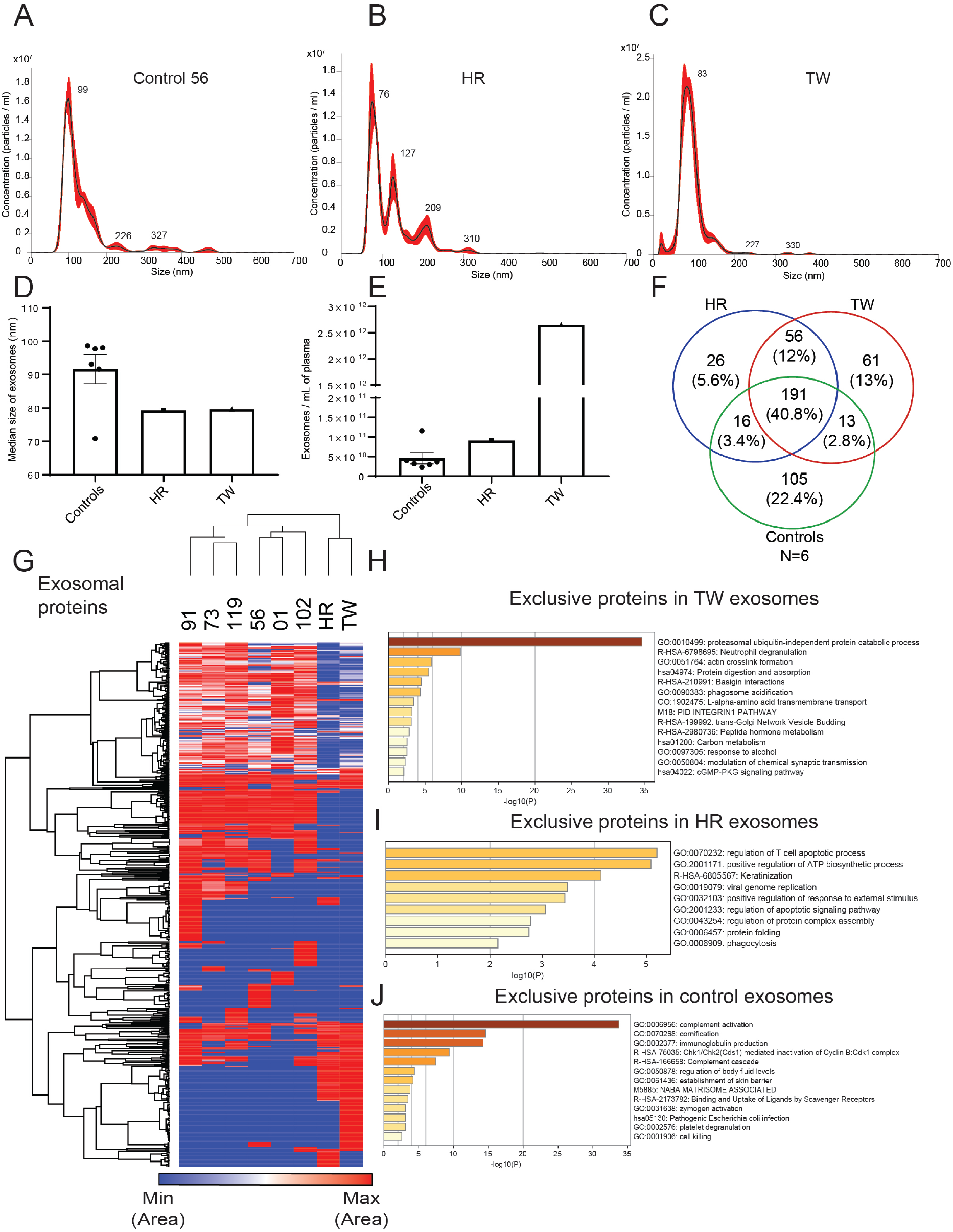
Characterization of plasma-derived exosomes isolated from HR and TW. Plasma samples were collected 3 years (TW) and 9 years (HR) post-flight. Nanosight profiles showing size distribution for exosomes isolated from the plasma of (A) Control, (B) HR, and (C) TW. Median size of exosomes (D) and exosome concentration (E) in TW (n=1), HR (n=1), and controls (n=6). (F) Venn diagram of exosomal proteins identified by mass spectrometry in plasma isolated from HR, TW and age-matched healthy controls. (G) Heatmap of plasma-derived exosomal proteins for HR, TW, and age-matched healthy controls. Pathway analysis of exclusive plasma-derived exosomal proteins from (H) TW, (I) HR, and (J) age-matched healthy controls.

Hierarchal clustering of the exosomal proteins revealed distinct signatures of HR and TW, which clustered apart from the six controls. Interestingly, classification of the pathways using Metascape (GO processes, KEGG pathways, Reactome gene sets, canonical pathways, and CORUM complexes)(Zhou et al., 2019) revealed that TW exosomes were enriched in proteins involved in proteasome pathways (**Fig. 6H**). TW exosomes also packaged CD14, a pro-inflammatory monocyte marker, consistent with the increase in CD14^+^ monocytes observed post-return to gravity in immune markers studied upon return to Earth (Gertz et al, 2020). Notably, basigin and integrin β1 proteins, which are correlated with cancer progression and inflammation (Hoshino et al., 2015, 2020; Keller et al., 2009; Yoshioka et al., 2014), were also detected in TW exosomes, but not in HR or healthy control exosomes.

Consistent with previous findings demonstrating microgravity downregulating adaptive immunity, particularly B cells (Cao et al., 2019), both TW and HR exosomes contained fewer immunoglobulins compared to healthy controls (**Fig. 6G**). Surprisingly, two brain-specific proteins, Brain-specific angiogenesis inhibitor 1-associated protein 2 (BAIAP2) and Brain-specific angiogenesis inhibitor 1-associated protein 2-like protein 1 (BAIAP2L1), were found in TW plasma-derived exosomes (**Supplemental Table 1, Supplemental Fig. 4A**), yet were not detected in the plasma of HR or healthy controls. In contrast, HR exosomal cargo was enriched in proteins associated with regulation of apoptotic pathways (Théry et al., 2001) and ATP biosynthesis (**Fig. 6I**). Moreover, we observed that the 20S proteasome, but not the regulatory 19S proteasome, is found uniquely associated with the plasma-circulating exosomes in the flight subject (TW) 3 years after his return to Earth (**Fig. 6G**). Finally, both TW and HR exosomes, but not controls, were enriched in specific components of the humoral immune response and leukocyte migration, including the CD53 tetraspanin (**Supplemental Table 2, Supplemental Fig. 4B**) which could reflect either biology shared by the twins or changes associated with travel to space; however, analysis of plasma exosome samples from genetically unrelated astronauts would be required to distinguish between these possibilities.

## Discussion

Our study focused on cfDNA and exosomes collected during the NASA Twins study, a longitudinal, multi-omic experiment examining the effects of long-term spaceflight on the human body. In particular, we revealed cf-mtDNA fraction to be a potential new biomarker of physiological stress during prolonged spaceflight, though the total cfDNA concentration is not significantly correlated with spaceflight. We further observed unique exosome and exosomal protein signatures within TW several years after the year-long mission, including an increased amount of exosomes and brain-specific proteins (BAIAP2 and BAIAP2L1). Of note, we identified multiple biases likely caused by ambient return (AR) blood draws from the ISS, including results of tissue of origin deconvolution through nucleosome positioning as well as cfDNA fragment length. As such, future studies will need to control for AR affects if they wish to examine these molecular dynamics. As an example, DNA could be extracted in space (Castro-Wallace *et al*., 2017) and either cryopreserved to increase its stable during transport or directly sequencing inflight to minimize biases and obtain results faster (McIntyre *et al*., 2016, McIntyre *et al*., 2019).

Interestingly, analysis of plasma exosomes isolated post-return to Earth revealed unique alterations in TW relative to HR and healthy controls, such as a dramatic increase in the number of circulating particles as well as changes in the types of protein cargo. Since the majority of plasma circulating exosomes are derived from immune cells, it is likely that these alterations reflect immune dysfunction associated with space travel and return to gravity. Specifically, the reduction in TW exosomal immunoglobulin levels and the presence of CD14, a macrophage marker, may signal a shift towards innate immunity, as even short-term chronic exposure to cosmic radiation and microgravity leads to a decrease in adaptive immune cells (Cao et al., 2019; Fernandez-Gonzalo et al., 2017). However, circulating exosomes also reflect systemic changes in homeostasis and physiology, as demonstrated by the packaging of brain-specific proteins in TW which were not seen in control exosomes, which may indicate long-term altered expression of exosomes from the brain after spaceflight. Previous studies had shown that microgravity affects tight junction protein localization within intestinal epithelial cells (Alvarez et al., 2019). It is conceivable that prolonged space travel could exert similar effects on tight junctions within the blood-brain barrier, allowing for more exosomes to enter the peripheral blood.

One remarkable finding of our study is that the 20S proteasome, but not the regulatory 19S proteasome, is found uniquely associated with the plasma-circulating exosomes in the flight subject 3 years post his return to Earth. Recent research has discovered the ubiquitin-independent proteolytic activity of the 20S proteasome and its role as the major degradation machinery under oxidizing conditions (Aiken et al., 2011; Deshmukh et al., 2019; Pickering and Davies, 2012).

Elevated levels of 20S proteasome have been detected in the blood plasma from patients with various blood cancers, solid tumors, autoimmune diseases and other non-malignant diseases (Deshmukh et al., 2019; Sixt and Dahlmann, 2008). It is also reported that active 20S proteasomes within apoptotic exosome-like vesicles can induce autoantibody production and accelerate organ rejection after transplantation (Dieudé et al., 2015), reduce the amount of oligomerized proteins (Schmidt et al., 2020).and reduce tissue damage after myocardial injury (Lai et al., 2012), and are correlated with cancer and other pathological status such as viral infection and vascular injury (Dieudé et al., 2015; Gunasekaran et al., 2020; Tugutova et al., 2019). The elevated circulating exosomal 20S proteasome in the flight subject may reflect the increased physiological need to clear these proteins resulting from long-term blood, immune or other physiological disorders caused by various stress factors during the flight or return to gravity (Ben-Nissan and Sharon, 2014, Vernice et al, 2020). Study of plasma exosomes obtained from flight subjects at other time points including pre- and inflight will be necessary to further examine whether plasma exosomal proteasome can serve as biomarker for pathological processes associated with space flight.

There are limitations in the study design that prevent broad biological conclusions. First, the sample number is too small to control for all types of potential biases and results may be somewhat driven by individual health issues. Second, there is no comparable experimental data to date and the effect of return to gravity in the Soyuz capsule on the integrity of the sampled material is unknown. Third, the exosome samples have been taken post-flight and can only inform about long-term effects of extended spaceflight. However, this study stands as a demonstration of the applications and possibilities of utilizing cfDNA and exosome profiling to monitor astronaut health and can improve the study design of future missions and research (Iosim et al, 2019, Nangle et al, 2020).

In summary, we identified cell-free mitochondrial DNA (cf-mtDNA) as a novel biomarker of physiological stress during prolonged spaceflight, which is stable even during transport from the ISS. However, we demonstrated that transport-induced biases for cell-type deconvolution from cfDNA needs to be improved in order to be used as a “molecular whole body scan”, and/or deployment of more real-time methods (e.g. inflight sequencing). Also, we observed that exosome concentration in plasma and unique exosomal proteins such as 20S proteasomes, CD14,and BAIAP2 demonstrate characteristic changes in the flight subject (TW), potentially caused by physiological stress during prolonged spaceflight. Overall, these data and methods provide novel metrics and data types that can be used in planning for future types of astronaut health monitoring, as well as help establish non-invasive molecular tools for tracking the impact of stress and spaceflight during future missions.

## Acknowledgements

We would like to thank the Epigenomics Core Facility and the Scientific Computing Unit (SCU) at Weill Cornell Medicine, as well as the Starr Cancer Consortium (I9-A9-071) and funding from the Irma T. Hirschl and Monique Weill-Caulier Charitable Trusts, Bert L and N Kuggie Vallee Foundation, the WorldQuant Foundation, The Pershing Square Sohn Cancer Research Alliance, NASA (NNX14AH51G (all Twins Study principal investigators); NNX14AB01G (S.M.B.); and NNX17AB26G (C.E.M.), NNX14AH52G), the National Institutes of Health (R25EB020393, R01NS076465, R01AI125416, R01ES021006, R01AI151059, 1R21AI129851, 1R01MH117406), TRISH (NNX16AO69A:0107, NNX16AO69A:0061, NIH/NCATS KL2-TR-002385), the Bill and Melinda Gates Foundation (OPP1151054), the Leukemia and Lymphoma Society (LLS) grants (LLS 9238-16, Mak, LLS-MCL-982, Chen-Kiang).

## Author Contributions

CEM, DB and DCL conceived the study, DB, CM, EA, SO, FAV, HZ, IM, KG and PB and, wrote the manuscript DB,BS, FEG, DBU, DPK, KHY, KN, TL, VR, FAV sample collection and/or processing DB, CM, SO, HZ, IM, JF, KG and PB Bioinformatic and Analytics, AM, BS, CEM, CM, CW, EA, FEG, IV, MC, MPS, RKB, SL, SO and TM review manuscript and guided interpretation. All authors read and approved the manuscript.

## Methods

### Sample collection

In the NASA Twin Study spanning 24 months we collected blood samples at 12 time points from the twin on earth (HR) and 11 time points from the twin in space (TW), as previously described(Garrett-Bakelman et al., 2019). From TW, samples were collected before the flight (PRE-FLIGHT), during the flight (FLIGHT) and after the flight (POST-FLIGHT). Specimens were processed as previously described(Garrett-Bakelman et al., 2019). Briefly, whole blood was collected in 4mL CPT vacutainers (BD Biosciences Cat # 362760,) per manufacturer’s recommendations, which contained 0.1M sodium citrate, a thixotropic polyester gel and a FICOLL Hypaque solution. Hence our specimens were not exposed to heparin. Samples were mixed by inversion. Samples collected on ISS were stored at 4°C after processing and returned by the Soyuz capsule. There was an average of 35-37 hours from collection to processing, including repatriation time. Plasma was obtained by centrifugation of the CPT vacutainers at 1800 X g for 20 minutes at room temperature, both for the ISS and for the ground-based samples. Finally, plasma was collected from the top layer in the CPT vacutainer and flash frozen prior to long term storage at - 80^°^C.

To simulate batch effects between fresh material (samples collected on earth) and ambient return material (samples collected during flight and returned via Soyuz capsule at 4°C), we generated 3 control samples (MS) representing fresh (FR) and ambient return (AR) material as described before (Garrett-Bakelman et al., 2019). Whole blood of a male volunteer of similar age and ethnicity as HR/TW was collected in three CPT tubes. Plasma was collected and stored as described for the TW and HR specimens. To generate the ambient return control (AR) two CPT vacutainers were shipped at ambient temperature (4°C) from Stanford University to Weill Cornell Medicine and back as air cargo. The returned CPT vacutainers were spun at 300 X g for 3 minutes and aliquoted. One aliquot from each tube was spun once more at 1800 X g for 3 minutes to completely clear the plasma of cell debris, resulting in the final AR controls. The aliquoted plasma was stored at -80C.

### cf DNA extraction and comparison of cfDNA concentrations in ground and flight subjects

Between 250ul and 1 ml plasma was retrieved from HR, TW and MS samples. The frozen plasma was thawed at 37C for 5min and spun at 16000g for 10 minutes at 4C to remove cryo-precipitates. The volume of each plasma sample was brought up to 1ml using sterile, nuclease-free 1X phosphate buffered saline pH 7.4. Circulating cell-free nucleic acid (ccfNA) was extracted using the Qiamp Circulating Nucleic Acid kit (Qiagen, USA) following the manufacturer’s protocol. ccfNA was extracted in 50ul AE buffer. Concentration and size distribution information was obtained by running 1ul of ccfNA on the Agilent Bioanalyzer using the High Sensitivity DNA chip (Agilent technologies, CA, USA). ∼15ul aliquots were set aside for cell-free DNA or DNA methylation analyses and stored at -80C. A range of extracted cfDNA of 1ng-38ng/mL plasma has been reported for healthy donors, while cancer patients often show higher levels of 30-50ng/ml (Table 1). In the HR, TW and control samples we extracted between 6.7 ng/ml and 79.9 ng/ml plasma (mean = 27.9 ng/ml, median = 23 ng/ml)(Table 1). We tested if there is a significant difference between HR, TW or MS as well as FR and AR samples using Wilcoxon rank test (R function wilcox) for pairwise comparisons and ANOVA (R function anova) for multi-group comparisons. We furthermore visualized the distributions of the groups HR, TW and MS as boxplots using the R package ggplot2.

### Q-PCR Analysis of cfDNA

The frozen plasma was thawed at 37C for 5min and spun at 16000g for 10 minutes at 4C to remove cryo-precipitates. DNA level in samples was measured by SYBR Green dye-based qPCR assay using a PRISM 7300 sequence detection system (Applied Biosystems) as described previously (Nakahira PLoS Med. 2013, Garrett-Bakelman Science 2019, PMIDs: 24391478 and 30975860). The primer sequences were as follows: human NADH dehydrogenase 1 gene (hu mtNd1): forward 5’-ATACCCATGGCCAACCTCCT-3’, reverse 5’-GGGCCTTTGCGTAGTTGTAT-3’. Plasmid DNA with complementary DNA sequences for human mtDNA was obtained from ORIGENE (SC101172). Concentrations were converted to copy number using the formula; mol/gram×molecules/mol = molecules/gram, via a DNA copy number calculator (http://cels.uri.edu/gsc/cndna.html; University of Rhode Island Genomics and Sequencing Center).

The thermal profile for detecting mtDNA was carried out as follows: an initiation step for 2 min at 50°C is followed by a first denaturation step for 10 min at 95°C and a further step consisting of 40 cycles for 15 s at 95°C and for 1 min at 60°C. MtDNA levels in all of the plasma analyses were expressed in copies per microliter of plasma based on the following calculation: c=Q x VDNA/VPCR x 1/Vext; where c is the concentration of DNA in plasma (copies/microliter plasma); Q is the quantity (copies) of DNA determined by the sequence detector in a PCR; VDNA is the total volume of plasma DNA solution obtained after extraction; VPCR is the volume of plasma DNA solution used for PCR; and Vext is the volume of plasma extracted.

### Library generation and sequencing

DNA libraries were generated using the NEBNext DNA Library Preparation Kit Ultra II (New England Biolabs, USA). Libraries were generated using 15ul of the ccfNA according to the manufacturer’s instruction. Following end-repair and dA-tailing, adaptor ligation was performed using 15-fold diluted adaptors. After removal of free adaptors using Agencourt magnetic beads (Beckman Coulter, USA), the libraries were PCR-amplified for 12 cycles using primers compatible Illumina dual-index sequences. Following bead cleanup for primer removal, the libraries were run on the Agilent Bioanalyzer to estimate size and concentration. All libraries were pooled at equal concentration and sent to New England Biolabs, Ipswich MA for sequencing. Preliminary sequencing on Illumina Miseq indicated the presence of adaptor dimers in some of the libraries. Therefore, the individual libraries were subjected to an additional round of bead purification and size and concentration estimation. Subsequently, all libraries were pooled again and sequenced on the NovaSeq 6000 using an S2 flow cell and 200-cycle kits (2×100). We finally obtained 4.9 and 4.1 billion reads passing quality filters.

### cfDNA sequence analysis

Samples were de-multiplexed using the standard Illumina tools. Low quality bases were trimmed and Illumina-specific sequences and low quality sequences were removed from the sequencing data using Trimmomatic-0.32. Filtered, paired-end reads were aligned using BWA-mem to the hg38 human reference genome with the bwa-postalt option to handle alternative alignments. The resulting BAM files were post-processed (e.g. sorted) using samtools. Duplicate sequences were removed and only reads aligning in concordant pairs were used for further analysis. The fragment length distribution was generated by plotting the distance between read 1 and 2 obtained from the BAM file of each sample. Histograms and boxplots of the fragment length distribution for the autosomes (Figure 1) and for the mitochondrial genome (Figure 2) for all samples were generated using the R package ggplot2.

### Analysis of cell free mitochondrial DNA

cfDNA read counts by chromosome (including the mitochondrial genome labeled ChrMT) were extracted using a ‘edtools coverage -a feature_file -b sample.bam –counts’, where feature_file contains the definition (name, start, end) of all chromosomes. Read counts per feature were length-normalized using the well-established reads-per-kilobase per million formula frequently applied to normalize RNA-seq data(Mortazavi et al., 2008). We used the R package ggplot 2 to visualize differences in the normalized cfDNA fraction (RPKM) originating from the mitochondrial genome between HR, TW and MS and over time during the mission (longitudinal analysis). We used Wilcoxon rank test (R function Wilcox) to test if the measurements of two conditions (e.g. TW on ground vs. TW in flight) are significantly different. To analyze the differences among multiple groups we applied ANOVA (R function anova).

### Nucleosome positioning analysis

Filtered, paired-end reads were aligned using BWA-mem to the hg37 human reference genome and post-processed using samtools: duplicate sequences were removed, and only reads aligning in concordant pairs were used for final analysis. The sequence read coverage in 10-kbp windows (−5 kbp to 5kbp) around the transcription start sites of all genes was determined using the samtools depth function. From the positions of the reads, nucleosome occupancy was inferred, and its periodograms calculated. A list of transcription start sites organized by transcriptional activity (measured in FPKM) was used to assign activity(Ulz et al., 2016). The depth of coverage was summed across genes according to transcriptional activity category (as depicted in Figure 4A). The coverage was normalized by subtracting the mean value from the intervals [TSS-3 kbp, TSS-1 kbp] and [TSS+1 kbp, TSS+3 kbp]. According to the method in Snyder et al., 2015, FFT values for the periods of 193-199bp were correlated with the gene level expression matrix, and the resulting tissue-periodicity correlations ranked by the value of Pearson’s correlation coefficient and clustered (Ward method with Euclidean distances) to investigate characteristics of tissue-of-origin dependent on sample type.

### Purification and Mass spectrometry analysis of plasma-circulating exosomes

Blood plasma was collected from TW 3 years post return of TW to Earth. Blood was also collected from HR (within one day of blood collection from TW) and from six age-matched healthy controls. Exosomes were purified by sequential ultracentrifugation, as previously described (Hoshino et al., 2015). Plasma samples were centrifuged for 10 minutes at 500xg, 20 minutes at 3,000xg, 20 minutes at 12,000xg, and the supernatant was collected and stored at -80°C for exosome isolation and characterization by NTA (NanoSight NS500, Malvern Instruments, equipped with a violet laser (405 nm). Samples were thawed on ice and centrifuged at 12,000xg for 20 min to remove large microvesicles. Exosomes were collected by spinning at 100,000xg for 70min, washed in PBS and pelleted again by ultracentrifugation in a 50.2 Ti rotor, Beckman Coulter Optima XE or XPE ultracentrifuge. The final exosome pellet was resuspended in PBS, and protein concentration was measured by BCA (Pierce, Thermo Fisher Scientific). Mass spectrometry analyses of exosomes were performed at the Rockefeller University Proteomics Resource Center using 10 μg of exosomal protein as described previously(Hoshino et al., 2015; Zhang et al., 2018). Heatmap and complete Euclidean clustering was performed with Morpheus, (https://software.broadinstitute.org/morpheus). Pathway analysis was performed with Metascape(Zhou et al., 2019).

## Figures, Tables, and Supplementary Tables/Figures for

**Supplemental Figure 1:**
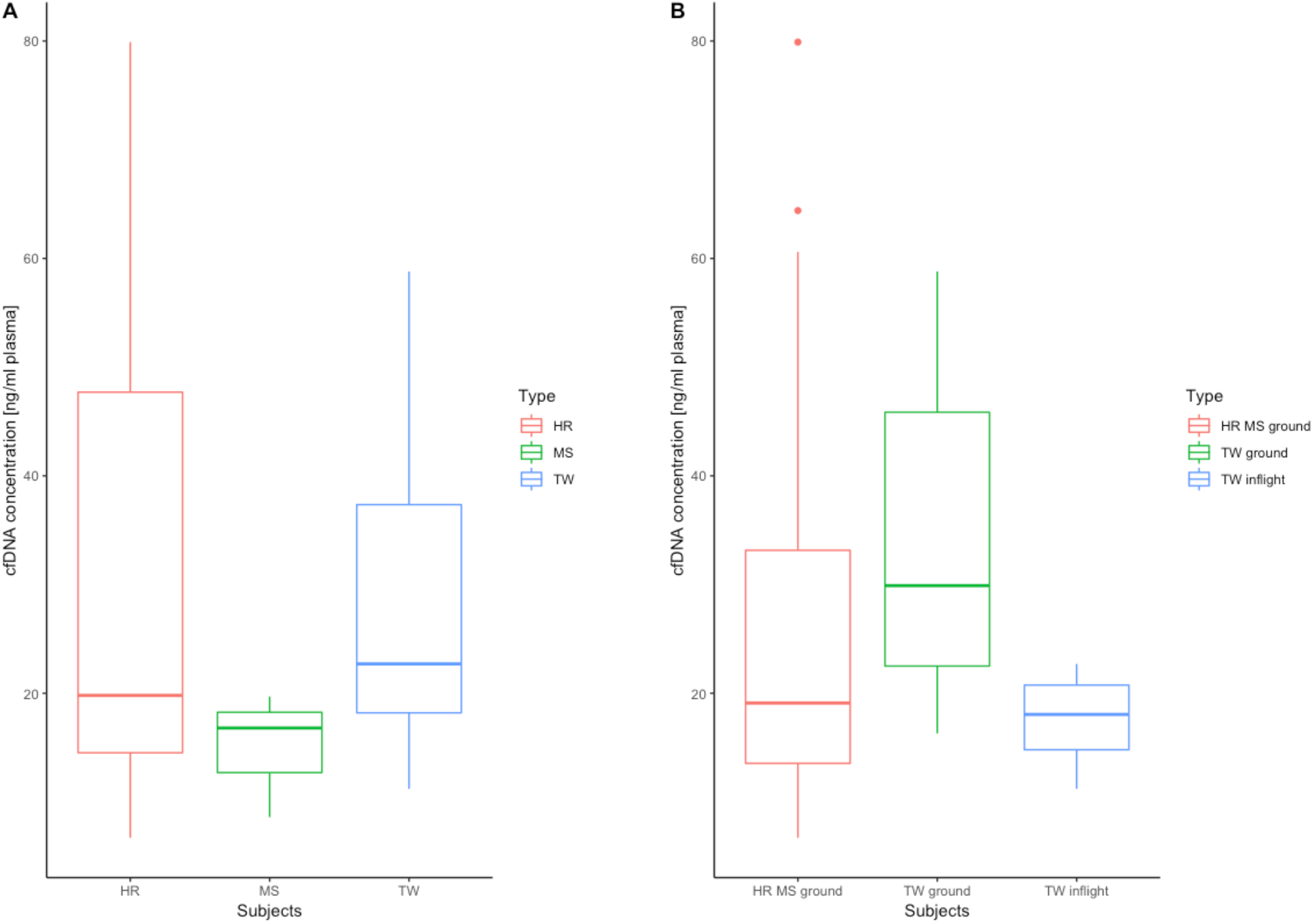
cfDNA concentration in plasma of ground subject (HR), flight subject (TW) and ground controls (MS). (A) Comparison of cfDNA concentrations between HR, MS and TW samples. (B) Comparison of TW pre- and –post-flight to TW inflight and to the combined ground samples of HR and MS. No significant differences were observed.

**Supplemental Figure 2.**
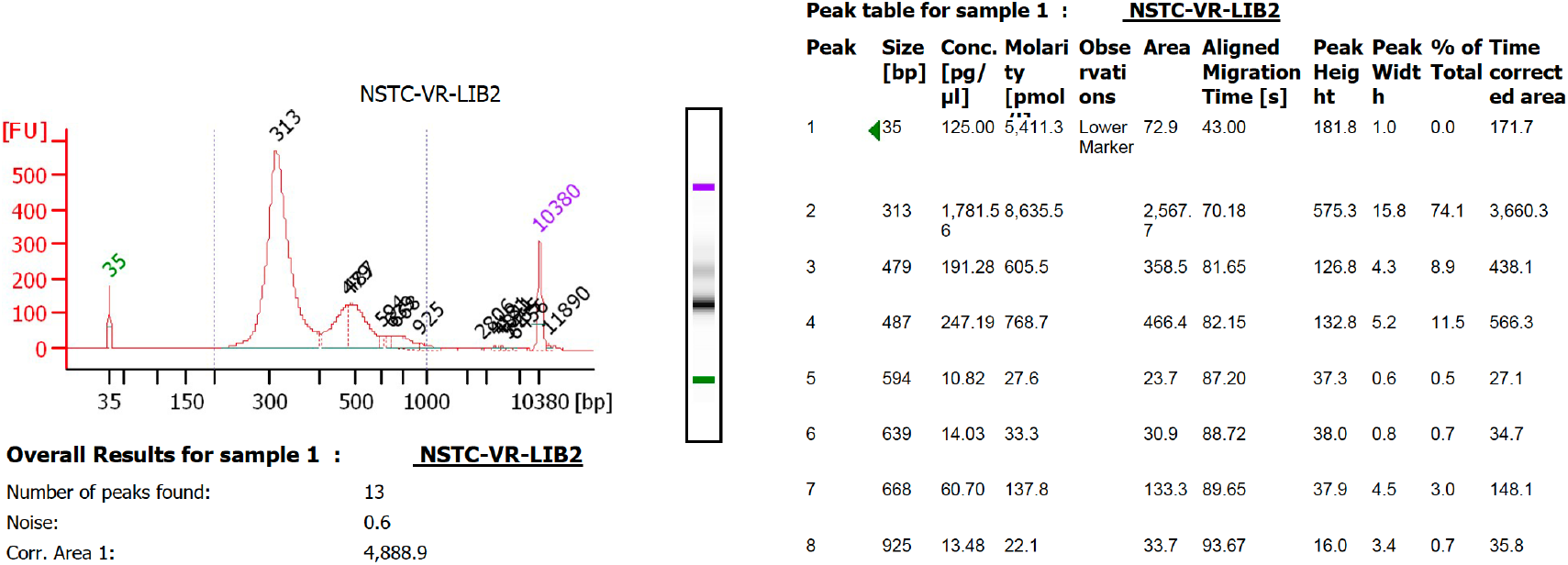
Library Fragment distribution for the cell-free DNA libraries. Pooled libraries were run on the Agilent Bioanalyzer 2100, with the entire fragment range area estimated to be 4,888.9. The first fraction peak was estimated to be at 313bp and the second peak was at 466bp. Given that the Illumina adapters add 120bp to each fragment size, this means that the estimated size of the first fragment set is 193bp and the second set is 346bp. The total area of the first peak represents 74.8% (3,660.3/4,888.9) of the signal.

**Supplemental Figure 3:**
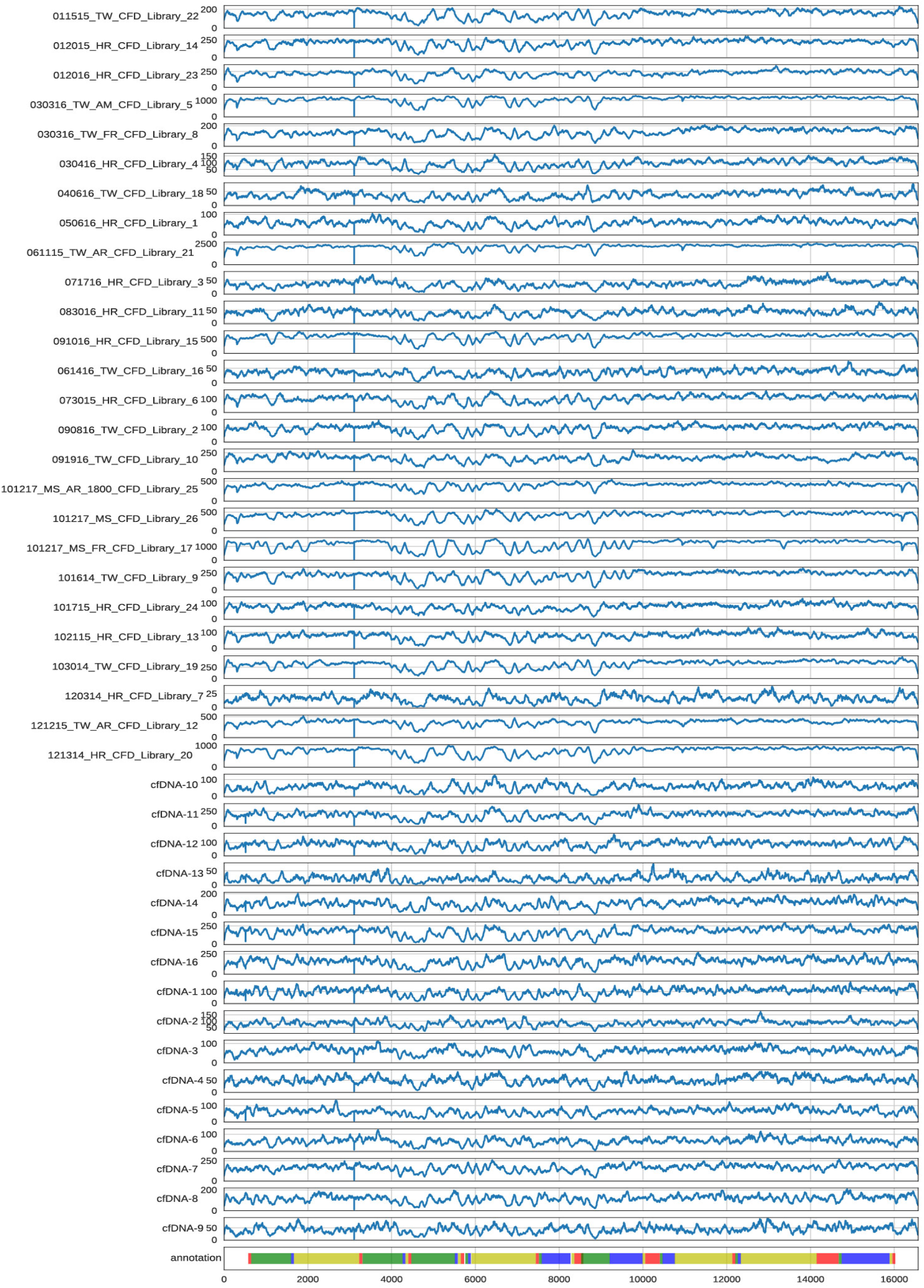
Read coverage distribution across the mitochondrial genome for all samples analyzed in this study. We observed continuous coverage of the complete mitochondrial genome in all samples.

**Supplemental Figure 4:**
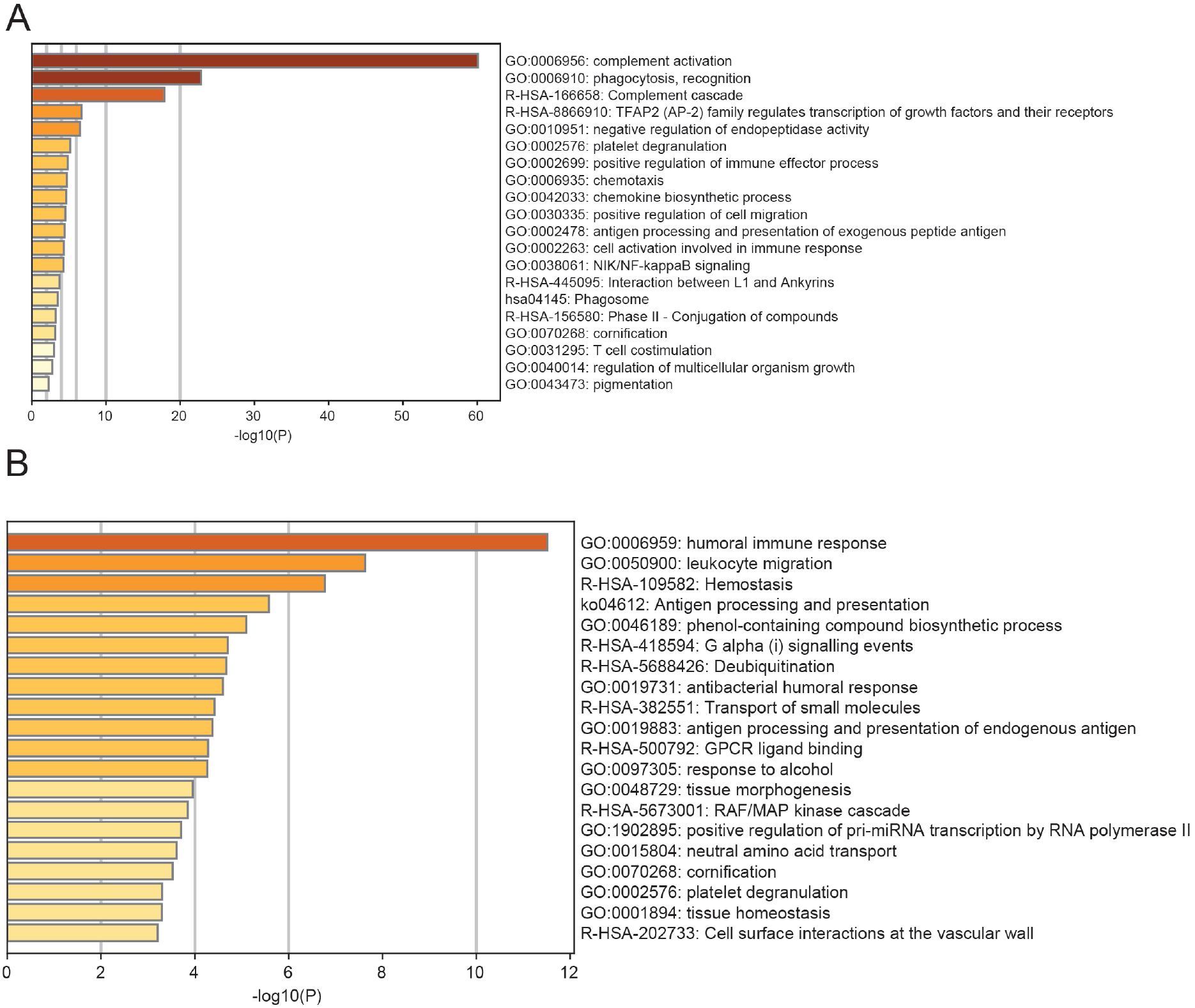
(A) Pathway analysis of inclusive plasma-derived exosomal proteins from TW (n=1), HR (n=2), and (J) age-matched healthy controls (n=6). (B) Pathway analysis of inclusive plasma-derived exosomal proteins from TW and HR excluding age-matched healthy controls

## Supplementary Tables

**Supplemental Table 1:**
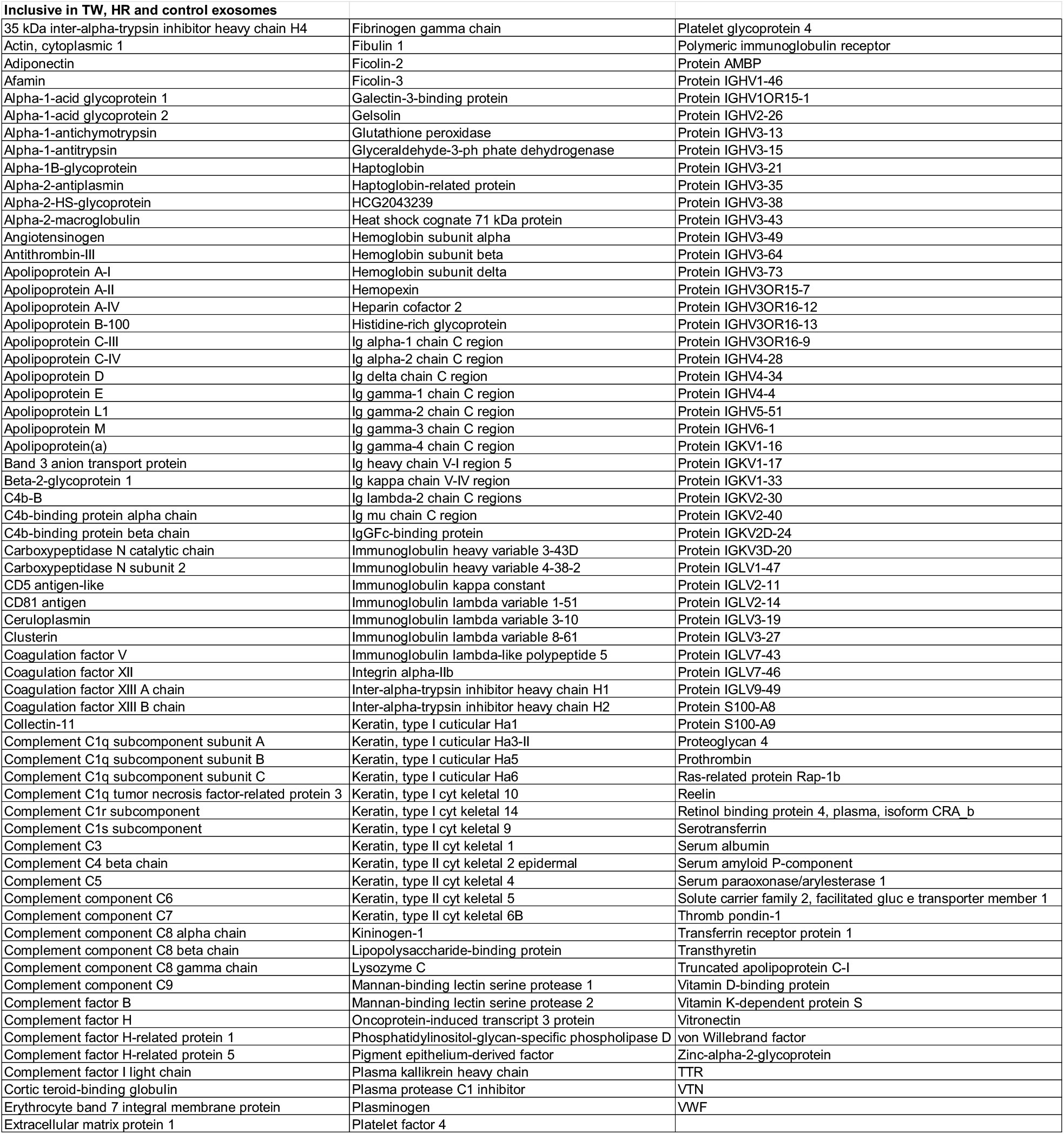
List of inclusive proteins in exosomes isolated from the plasma of TW, HR and age-matched healthy controls.

**Supplemental Table 2:**
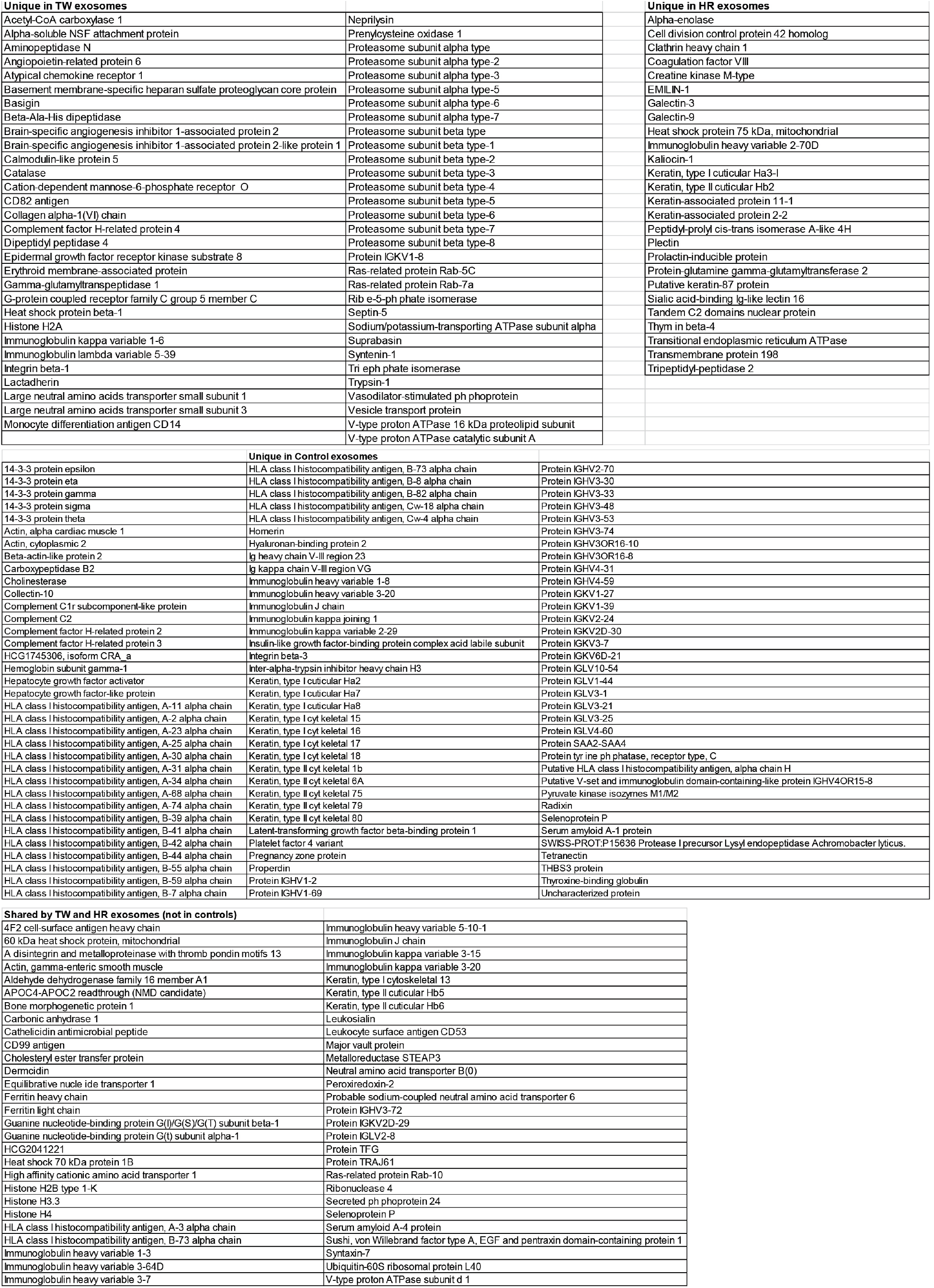
List of unique proteins in exosomes isolated from the plasma of TW, HR and age-matched healthy controls.

